# Higher-order interdependencies of the epileptogenic network in presurgical resting-state fMRI

**DOI:** 10.64898/2026.09.24.753209

**Authors:** Jacob Knight, Matteo Neri, Julia Makhalova, Hugo Dary, Samuel Medina-Villalon, Jean-Philippe Ranjeva, Andrea Brovelli, Fabrice Bartolomei, Maxime Guye, Roy AM Haast

**Affiliations:** Aix-Marseille Université, CRMBM, CNRS UMR 7339, Marseille, France; CEMEREM, APHM, Hôpital de la Timone, Marseille, France; Service d’épileptologie et de rythmologie cérébrale, APHM, Hôpital de la Timone, Marseille, France; INS, INSERM UMR 1106, Aix-Marseille Université, Marseille, France; Institut de Neurosciences de la Timone UMR 7289, Aix Marseille Université, CNRS, 13005, Marseille, France

## Abstract

Drug-resistant focal epilepsy (DRFE) is understood as a complex brain network disorder. As a result, conventional pairwise functional connectivity may fail to capture the higher-order interactions between groups of epileptogenic regions. We applied multivariate information theory to stereotactic electroencephalography-defined networks in retrospective resting-state fMRI analysis to determine whether the overall strength and dominant type of interdependencies between three or more regions, represented by the S-information and O-information respectively, could offer clinically relevant insights prior to surgical intervention in DRFE. We found networks involving the epileptogenic zone to be mostly dominated by redundant information-sharing, which was predicted by stronger structural and functional connections between regions. Non-temporal lobe epilepsy patients demonstrated greater redundancy and stronger comparative interdependencies than temporal lobe epilepsy patients, in line with a more complex network architecture. Among non-temporal lobe cases with postoperative follow-up, not seizure-free patients displayed more positive O-information deviation scores than seizure-free patients, suggesting elevated redundancy in networks associated with poorer surgical outcomes. Our findings potentiate the S-information and O-information as complementary higher-order descriptors of the epileptogenic network within personalised presurgical assessments.

## Introduction

Network neuroscience offers a framework for understanding the epileptic brain as a perturbed complex system. Proposing a non-localised area of onset, the epileptogenic network model effectively demonstrates how seizures are accommodated in patients presenting with drug-resistant focal epilepsy (DRFE) prior to potential surgery^1-3^. Derived from intracranial recordings, the epileptologist establishes a hierarchical organisation of nodes that contribute to seizure activity, with electrode contacts typically assigned to either the epileptogenic zone network (EZN) or propagator zone network (PZN)^2,4^. In stereotactic electroencephalography (SEEG), quantified methods have been proposed to more objectively classify involved networks. The epileptogenicity index (EI) offers a spectrotemporal method for elucidating the EZN and PZN, with higher values denoting structures involved early in the ictal process^5,6^. Subsequent surgical removal of the whole or part of the EZN disrupts seizure generation, a practice most often performed in drug-resistant temporal lobe epilepsy (TLE) where it has proven a reliable treatment^7-9^.

Despite some improvement in recent years, achieving seizure freedom in patients with drug-resistant non-temporal lobe epilepsy (NTLE) remains difficult^10-12^. NTLE poses distinct challenges to the epileptologist, who must identify an often diffuse pathological system of structurally and functionally independent nodes^11,13,14^. Differentiating spared regions from the resection area holds particular importance in NTLE, with the epileptogenic zone often neighbouring or encompassing parts of the eloquent cortex^13,15^. Arising from numerous possible sites, the spatial arrangement of epileptogenic networks in NTLE is inevitably heterogeneous and often coincides with widespread disruptions to brain connectivity^16-18^. Ultimately, surgery tends to yield poorer outcomes in NTLE compared to patients with TLE, with about 25–65% considered seizure-free after extratemporal resections versus roughly 55–80% after temporal resection^12,19^. Characterising and comparing the properties of SEEG-defined TLE and NTLE networks in resting-state fMRI (rs-fMRI) could serve to bridge the gap in surgical efficacy between the two patient groups.

Previous rs-fMRI studies in focal epilepsy have primarily focused on bivariate temporal correlations between brain regions, commonly known as functional connectivity (FC), to localise areas of seizure onset and evaluate canonical resting-state networks^20-22^. With inconsistent results and non-concordance between rs-fMRI and SEEG-derived FC^23,24^, the study of DRFE offers an important opportunity to validate non-invasive imaging and alternative network models^25^. Recent evidence suggests that pairwise interactions alone may be insufficient in capturing emergent properties of networks^26-28^. Conceiving neural activity as the flow of information, multivariate information theory offers a powerful entropy-based framework for illustrating complex systems^28^. As a higher-order model, multivariate information theory can describe the collective dynamics of expanded brain networks comprising three or more regions from time series data^30,31^. Two complementary measures defined by Rosas *et al*.^32^ are particularly telling: the S-information, which quantifies the overall statistical dependence between nodes in a network and relates closely to measures of complexity, and the O-information, which characterises whether those interdependencies are predominantly redundant or synergistic. Opposing synergistic interactions where information is combined, redundant interactions reflect the sharing of copied or undifferentiated information between regions and support intramodular connections in the brain^33-35^. Whilst multivariate measures of redundancy have been applied to the aging brain^36,37^ and neurodegenerative disease^38^ in rs-fMRI, investigations in a specific medical setting remain underdeveloped. Animal models of TLE have shown perturbed information-sharing during the interictal resting-state and increased network redundancy in the early stages of epileptogenesis^38,39^. Given the abnormal synchronisation of distant structures observed during intracranial seizure recordings^2^, a conceivable link may exist between redundant information and pathological network activity in focal epilepsy.

Offering global descriptions of large-scale network organisation which may be missed by pairwise approaches, measures of multivariate information could provide further insight into the functional architecture of groups of brain regions involved in focal seizures prior to surgical intervention. We hypothesise that epileptogenic networks comprising three or more regions are dominated by redundant information-sharing during the interictal resting-state and that the S- and O-information capture higher-order interactions that distinguish TLE and NTLE networks from non-epileptic systems. To this end, we presently define the higher-order information of SEEG-defined epileptogenic networks from the presurgical rs-fMRI data of patients with DRFE (see Fig. 1). We contextualise the synergy-redundancy balance of these networks by quantifying their relationship to conventional structural and functional MRI-based connectivity measures. Finally, to explore biomarker potential, we compare the higher-order information of each epileptogenic network to patient-level null and matched healthy control distributions, before discussing associations to surgical outcome. Taken together, this work provides a first multi-modal investigation into the clinical utility of multivariate information theory in focal epilepsy, establishing a foundation for broader exploration of the epileptogenic network’s higher-order properties.

**Fig. 1:**
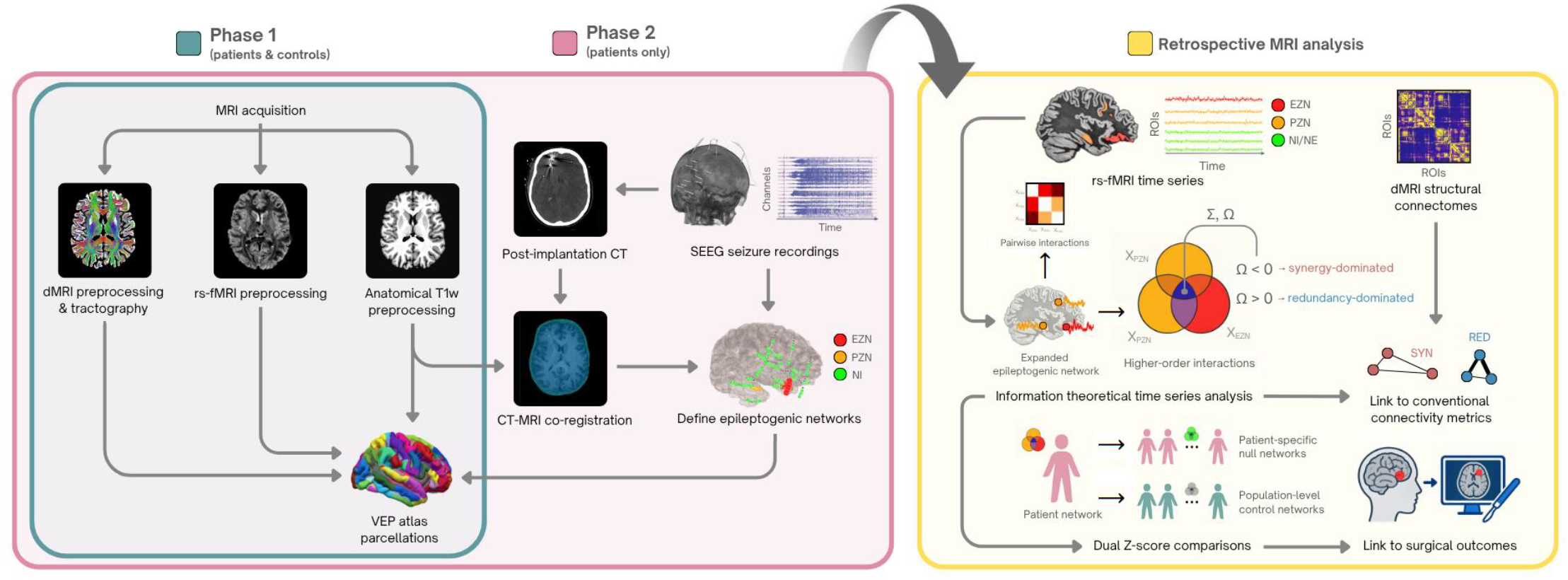
Overview of presurgical workup for MRI-based analysis of patient epileptogenic and propagator zone networks seeded from intracranial seizure recordings. Patient MRI was performed as part of Phase 1 presurgical workup and included anatomical T_1_-weighted (T1w), resting-state functional (rs-fMRI) and diffusion-weighted (dMRI) acquisitions. This was followed by Phase 2 stereotactic electroencephalography (SEEG) from which epileptogenic zone (EZN) and propagator zone (PZN) network membership or non-involved (NI) status was assigned to contacted regions-of-interest (ROIs) as per Virtual Epileptic Patient (VEP) atlas parcellations. The network-level S-information (Σ), O-information (Ω) and functional connectivity were calculated from rs-fMRI time series data and structural connectivity computed from dMRI tractography data in retrospective MRI analysis for each epileptogenic network. Patient networks were then compared to healthy controls (grey) and patient-specific null networks sampled from NI and non-explored (NE) regions (bright green) before exploring relationships to postoperative seizure freedom.

## Methods

### Participants

We studied a cohort of patients (n = 66) diagnosed with DRFE who underwent presurgical evaluation at our facility from June 2017 to October 2023. Seven Tesla (7T) MRI scans were collected within the scope of two prospective studies, namely the RHU EPINOV clinical trial (NCT03643016) and NEURO7T (2016-A01785-46). Following official French guidelines^40^, intracranial electrophysiological recordings were systematically performed for all patients as part of routine presurgical workup. We also recruited a population of healthy controls (n = 42) with no history of neurological disorder who underwent a 7T MRI scan between July 2017 to September 2023 for Z-score analysis. Each study received separate approval from the local ethics committee (*Comité de Protection des Personnes sud Méditerranée 1*) and all participants provided written informed consent in compliance with the Declaration of Helsinki. The current investigation complies with confidentiality requirements set by the national personal data regulation committee (*Commission Nationale de l’Informatique et des Libertés*).

### 7T MRI acquisition

All patients and healthy controls underwent 7T MRI B_1_^+^ and T^1^ mapping followed by rs-fMRI. Scans were conducted by a whole-body Magnetom or Magnetom Terra system (Siemens Healthineers, Erlangen, Germany) using a ^1^H 1Tx/32Rx head coil (Nova Medical, Wilmington, USA).

Ultra-high resolution B_1_^+^ and T_1_-weighted (T_1_w) maps with an isotropic voxel size of (0.6 mm)^3^ were acquired using a spin echo-based sequence that assessed the ratio of consecutive spin and stimulated echoes^41^ and the 3D Magnetization Prepared with 2 Rapid Acquisition Gradient Echoes (MP2RAGE) respectively^42^. Blood oxygen level-dependent (BOLD) rs-fMRI data with an isotropic nominal voxel size of (1.6 mm)^3^ was acquired using 2D Multi-Band (MB) Echo Planar Imaging (EPI) sequences (MB factor = 5, in-plane acceleration factor = 2, TR/TE_MAGNETOM_ = 1000/22 ms, TR/TE_TERRA_ = 1080/25.2 ms, α = 38°, partial Fourier = 0.88, TA_MAGNETOM_ ≈ 3min, TA_TERRA_ ≈ 4 min) yielding 85 slices across 200–220 volumes. Additional rs-fMRI data was acquired in the reverse phase-encoding (PE) direction (left-right to right-left) with otherwise identical scanning parameters to correct for EPI-related geometric distortions.

Amongst patients, diffusion MRI (dMRI) images with a (1.1 mm)^3^ voxel size were attained using single-shot EPI diffusion-weighted sequences (80 encoding directions, TR/TE_MAGNETOM_ = 6000/79.2 ms, TR/TE_TERRA_ = 5500/78.4 ms, b = 0, 2000 s/mm2, α_MAGNETOM_ = 90°, α_TERRA_ = 70°, partial Fourier = 0.75, TA ≈ 10 min) delivering 120 slices across 93 volumes acquired twice in both PE directions (anterior-posterior to posterior-anterior). All rs-fMRI and diffusion-weighted images were visually inspected for artifacts and susceptibility-induced signal loss. For each rs-fMRI run, whole-brain frame displacement (FD) and temporal signal-to-noise ratio (tSNR) were extracted using MRIQC^43^ (v24.0.2) for post-hoc quality control (see Supplementary Fig. 1B).

### Intracranial electrophysiological recordings

SEEG was performed using intracerebral multi-contact ALCIS® electrodes (10-18 contacts of 2 mm length, 0.8 mm diameter and 1.5 mm spacing) implanted using a frameless ROSA ONE® stereotactic surgical assistant. Recordings of spontaneous seizures and wakeful resting-state periods were acquired for each patient. Signals were recorded on a 256-channel NATUS® system and sampled at 512 Hz to a hard disk (16 bits/sample) with no digital filter. Acquisition included high-pass (0.16 Hz at -3 dB) and anti-aliasing low-pass (170 Hz at 512 Hz) filters. Post-implantation, each patient underwent a cranial computed tomography (CT) scan to verify implantation accuracy and the absence of complications.

### Anatomical image preprocessing

T_1_w images were skull-stripped and B_1_-corrected using PreSurfer^44^ (v1.1.0) before intensity non-uniformity correction with *N4BiasFieldCorrection*^45^ implemented in ANTs^46^ (v2.6.0). Tissue segmentation distinguishing cerebrospinal fluid (CSF), white-matter (WM) and grey-matter (GM) was performed on brain-extracted T_1_w images using *fast* from FSL^47^ (v6.2.7). Brain surface reconstruction was performed using *recon-all* from FreeSurfer^48^ (v8.1.0). Volume-based spatial normalisation to standard Montreal Neurological Institute (MNI) space with the *MNI152NLin2009cAsym* template was accomplished through non-linear registration using ANTs’ *antsRegistration*. Preprocessed anatomical images served as T_1_w references for the remainder of the preprocessing workflow.

### rs-fMRI time series preprocessing

Preprocessed BOLD images for patients and healthy controls were obtained using fMRIPrep^49^ (v25.1.2) which is based on Nipype^50^ (v1.10.0). Using a custom methodology, a reference volume was generated for head motion correction for each of the two rs-fMRI runs. A B_0_ fieldmap was estimated based on the two EPI references with opposing PE directions using AFNI’s (v16.2.07) *3dQwarp*^51^. From the estimated susceptibility distortion, a corrected EPI reference was calculated for co-registration with the anatomical T_1_w reference using FreeSurfer’s *bbregister* with six degrees of freedom^52^.

Head-motion parameters with respect to the BOLD reference images (transformation matrices, six corresponding rotation and translation parameters) were estimated before spatiotemporal filtering using FSL’s *mcflirt*^53^. The BOLD time series (including slice-timing correction) were then resampled to their original native space by applying a single composite transform using ANTs’ (v2.3.3) *antsApplyTransforms* and configured with Lanczos interpolation^54^ to correct for head-motion and susceptibility distortions. We will refer to these BOLD time series as preprocessed BOLD data.

Several confounding time series were calculated based on the preprocessed BOLD. Three region-wise global signals were calculated within the anatomically derived eroded CSF, WM and whole-brain (global signal) masks. The head-motion estimates calculated in the correction step were also placed within the corresponding confounds file. Six head-motion estimates (three translational, three rotational) and the three physiological signals (CSF, WM and global signal) were extracted from each file and used for confound regression before parcellation and temporal filtering. The choice for global signal regression was made in line with methodology from Varley *et al*.^30^.

The Virtual Epileptic Patient^55^ (VEP; https://ins-amu.fr/vep-atlas) atlas was computed from each subject’s FreeSurfer output and registered to corresponding preprocessed BOLD data. The BOLD time series was then parcellated using the VEP atlas and subjected to temporal filtering (high-pass: 0.009 Hz, low-pass: 0.08 Hz), detrending and standardisation using Nilearn’s (v0.13.1) *NiftiLabelsMasker*. To account for acquisition variability between scanners and non-steady-state signal, time series data was truncated to 200 timepoints and the first five volumes discarded. Finally, atlas labels were mapped to parcellated regions using the VEP atlas lookup table and excluded regions removed, delivering a 162 (region) × 195 (timepoint) labelled time series for each of the two rs-fMRI runs per subject.

### Structural connectomes

Diffusion image preprocessing and whole-brain tractography were completed amongst patients with quality data following visual inspection. We applied denoising, Gibb’s artifact removal, N4 bias field inhomogeneity correction and susceptibility distortion correction with FSL’s *topup*^56^ using two diffusion-weighted images in opposing PE directions per patient.

Following Eddy current and head-motion correction, resulting mean b = 0 images were used to register subject T_1_w maps to diffusion space using FSL’s *flirt*^53^ with six degrees of freedom and normalised mutual information cost. The previously computed VEP atlases were also registered to subject diffusion space.

Anatomically constrained tractography (ACT) was performed with MRtrix3 (v3.0.8). Following estimation of WM fibre orientation direction with constrained spherical deconvolution^57^, SIFT-based ACT^58^ was carried out to deliver 10 million streamlines. Resulting .*tck* files underwent two separate VEP atlas-based parcellations with *tck2connectome* to compute one streamline count-weighted and one streamline length-weighted 162 × 162 structural connectome per patient. Streamline count-weighted connectomes were normalised by regional node volume.

### Epileptogenic networks

Excluding artifacts and non-GM electrode contacts, signals from two seizure SEEG epochs per patient were investigated using AnyWave software^59^ (https://gitlab-dynamap.timone.univ-amu.fr/anywave). For each bipolar SEEG contact pair, the EI was calculated and used to define epileptogenic network partitions. The EI combines the ratio of fast frequencies (12.4– 97 Hz) relative to slower frequencies and the time of involvement of each electrode contact^5^. As per previous work^60,61^, derivations with an EI ≥ 0.4 and 0.1 ≤ EI < 0.4 were assigned to the EZN and PZN respectively. Contact pairs that displayed an EI ≤ 0.1 were considered to minimally contribute to epileptiform discharges and were assigned non-involved (NI) status.

For each patient, pre-implantation T_1_w MRI images were co-registered to post-implantation cranial CT images followed by automated anatomical localisation and labelling of each electrode contact using the GARDEL software from EpiTools^62^ (https://meg.univ-amu.fr/wiki/GARDEL). Co-registration with the VEP atlas was then performed for regional parcellation followed by the transformation of electrode coordinates (defined by a 5 mm sphere around each bipolar derivation^61^) to patient 7T MRI space with linear co-registration between CT, MP2RAGE and MNI template images. Electrode coordinates were then assigned to regions-of-interest (ROIs) as defined by the VEP atlas. When multiple electrodes contacted the same ROI, the most epileptogenic classification (EZN > PZN > NI) was assigned to that region.

Corresponding EZN, PZN and NI labels were mapped to parcellated VEP atlas regions of preprocessed BOLD rs-fMRI time series and dMRI weighted connectome data. ROIs not covered by SEEG were labelled as non-explored (NE). We will refer to patient ROIs assigned to the EZN or PZN as collectively belonging to the expanded epileptogenic network (EEN).

### Information theoretical analysis of rs-fMRI time series

We selected two measures of multivariate information defined by Rosas *et al*.^32^ for time series analysis: the S-information and the O-information, which are both derived from the total correlation^63^ and dual total correlation.^64^ Measures were calculated in binary digits for each epileptogenic network from preprocessed BOLD data using the HOI toolbox^65^ (https://brainets.github.io/hoi/). We together refer to the S-information and O-information as the higher-order information.

The total correlation (*TC*) and dual total correlation (*DTC*) are given by the following formulas:

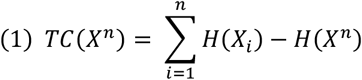

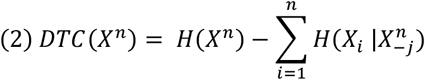

where *X*^*n*^ is a vector of *n* random variables and *H* represents the Shannon entropy. In equation 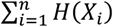 quantifies the amount of information carried by individual nodes in a network whilst *H*(*X*^*n*^) contains the information carried by individual nodes and their interactions. Upon subtracting the two quantities, we isolate the information carried by the network’s interactions alone; that is, the *TC*. We can consider the *TC* as a heuristic measure of redundancy. In equation (2), *H*(*X*^*n*^) once again represents the total information contained in the full network, including both individual nodes and their interactions. We then subtract the term 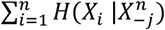 which captures the information that each node uniquely contains on its own; that is, the information that remains when we know the state of all other nodes. By conditioning on the rest of the network in this way, we isolate each node’s unshared entropy and remove it from the total network entropy to deliver the *DTC*. We therefore consider the *DTC* a heuristic measure of a synergy.

The S-information (Σ) and O-information (Ω) are constructed from the sums and differences of the *TC* and *DTC* respectively as shown in equations (3) and (4).

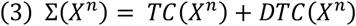

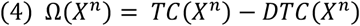

When the O-information is positive for a given network (Ω > 0), that network is dominated by redundant information. When negative (Ω < 0), the network is dominated by synergistic information. Importantly, we chose to compute these metrics at the network-level, which is defined as the maximal interaction order for the network in question^66^. For instance, a patient displaying an EZN where *N* = 3 and a PZN where *N* = 4 will have the S- and O-information calculated at orders 3, 4 and 7 for their EZN, PZN and EEN respectively. Details regarding this decision are examined in the Discussion.

### Connectivity analysis

Structural connectivity metrics were derived from whole-brain streamline-weighted connectomes. Metric selection was inspired by the structure-function coupling analysis performed by Liu *et al*.^67^. For each epileptogenic network, we extracted the following: mean connection weight (the average streamline count between regions), connection density (proportion of possible connections present), mean connection length (the average streamline path length between regions in millimetres), and network modularity (degree to which the network segregates into community structure). Whole-brain modularity was computed using NetworkX’s (v3.6.1) greedy modularity optimization algorithm^68^ followed by calculation of the largest module share for each network (the maximal number of regions falling into a single module divided by *N*).

Spatiotemporal properties of each epileptogenic network were extracted from rs-fMRI and anatomical data during computation of null distributions. Mean centroid Euclidean distance (millimetres) was computed from VEP atlas region centroid coordinates in MNI space. Mean FC was calculated from the Pearson correlation between BOLD signals for each node pair within a network and is reported as the average between the two rs-fMRI acquisitions.

### Z-score analysis

We obtained pairs of Z-scores from distinct reference distributions to answer two separate but related questions: (1) how does the higher-order information of patients compare to healthy controls and (2) how do those same interdependencies compare to networks of patient-specific NI/NE regions? Addressing (1), between-subject Z-scores (Z_b_-scores) were calculated by comparing the S- and O-information of each epileptogenic network against a distribution of homologous control networks. Each healthy control subject was assigned a network based on their rs-fMRI time series for the same VEP atlas regions forming the patient’s epileptogenic network. The S- and O-information were then computed for all homologue control networks, yielding a distribution (n = 42) of healthy control values per metric per patient seed for Z_b_-score calculation. Addressing (2), within-subject Z-scores (Z_w_-scores) were computed by comparing each epileptogenic network against a null distribution (n = 1000) extracted from regions of the patient brain not belonging to the EEN. Null networks were thereby randomly sampled from rs-fMRI time series data based on VEP atlas regions labelled as either NI or NE. Networks accepted into the null distribution were required to match the following criteria: (i) number of nodes (*N* equal to that of the epileptogenic network), (ii) node spatial arrangement (≤ 20mm difference in mean centroid Euclidean distance from that of the epileptogenic network) and (iii) node temporal synchronisation (≤ 0.1 difference in mean FC from that of the epileptogenic network). The S- and O-information were then computed for each matched network and added to the null distribution. Combinations failing to meet spatiotemporal constraints were rejected and resampling continued until 1000 null values per metric were obtained. This process was repeated with all homologous control networks for paired comparisons, with resulting Z_w_-scores averaged per metric per patient seed. Any networks where imposed spatiotemporal constraints made it computationally infeasible to obtain the target null distribution were excluded from further analysis. A sensitivity analysis involving a random subset of patients was also performed to test the effect of different null acceptance ranges on Z_w_-scores and sampling entropy (see Supplementary Fig. 3).

Z_b_- and Z_w_-scores were defined as the number of standard deviations from the respected distribution mean of each metric as per the following formula:

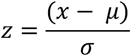

where *x* is the normalised S- or O-information for that network, *μ* is the comparative reference distribution mean and *σ* is the standard deviation for that distribution.

### Statistical analysis

Reported S- and O-information values for each network were normalised by the number of nodes (metric / *N*) following general approaches to accounting for dimensionality in multivariate information theory^69^. Resulting values were then averaged between the two rs-fMRI acquisitions. Reported Z_b_- and Z_w_-scores were derived from non-normalised metrics separately for each run before averaging. We performed statistical testing and multiple comparisons corrections using Pingouin (v0.6.1). Pearson’s r was calculated for each two-factor correlation analysis. Normality assumptions for all Z-score reference distributions were assessed via histogram inspection. We determined differences between group (patient vs control), epileptogenic network partition (EZN vs PZN vs EEN), focal epilepsy class (TLE vs NTLE), synergy-redundancy dominance (RED vs SYN) and surgical outcome (SF vs NSF) using parametric pairwise comparisons (paired and unpaired two-tailed t-tests; mean ± standard deviation and degrees of freedom reported). Abbreviations for test groups are summarised in Supplementary Table 2. Within-subject differences between network partitions were additionally assessed using the Friedman test. Because individual patients often contributed to more than one partition (with the EZN, PZN and EEN as related constructs), our observations were not fully independent at the subject-level. Accordingly, unpaired t-tests should be interpreted as network-level comparisons, with potential within-subject dependence acknowledged. Binomial tests from SciPy (v1.16.3) were used to assess proportions of synergy- and redundancy-dominated networks. In the case of multiple comparisons, p-values were corrected (p_corr_) per test following the Benjamini-Hochberg false discovery rate (FDR) procedure. PLS regression was carried out using scikit-learn (v1.5.2) to identify which combination of connectivity metrics best predicted O-information values. The optimal number of components was selected via five-fold cross-validation and statistical significance determined through permutation testing (n = 5000).

## Results

### Study cohort

Patient rs-fMRI and dMRI data was inspected for missing elements, acquisition artifacts and anatomical abnormalities. Combined SEEG and CT outcomes were inspected for epileptogenic network localisation. A total of eight patients were evicted from the study due to missing or corrupt imaging data. Two subsequent patients were removed due to motion artifacts and a further two due to morphological brain abnormalities. Finally, three patients displayed an EEN comprising only two parcellated regions and were deemed ineligible for higher-order analysis. A final study cohort of n = 51 patients each containing a unique configuration of epileptogenic networks was established (see Table 1), with most patients (n = 39) displaying an EZN, PZN and EEN where *N* ≥ 3. A subset of the final patient cohort possessed quality dMRI data (n = 42). No healthy control rs-fMRI data was discarded, and all subjects were utilised for Z-score analysis. Post-hoc analysis revealed little to no difference in the rs-fMRI results of each scanner (see Supplementary Fig. 1A).

**Table 1:** Demographics and clinical data summary of patients with drug-resistant focal epilepsy and healthy controls.

|  |  | Patients (n = 51) | Healthy controls (n = 42) |
| --- | --- | --- | --- |
| **Demographics** |  |  |  |
| Age, mean ± SD [median, min-max] | | 30.6 ± 12.4 [26, 15-57] | 33.5 ± 11.9 [28, 21-65] |
| Sex, n (%) | Male | 28 (54.9) | 19 (45.2) |
|  | Female | 23 (45.1) | 23 (54.8) |
| **Clinical descriptors** |  |  |  |
| Disease duration, mean ± SD [median, min-max] | | 16.6 ± 9.5 [14, 5-42] | |
| MRI lesion status, n | Positive | 22 |  |
|  | Negative | 29 |  |
| Epileptogenic zone lateralisation, n | Left | 22 |  |
|  | Right | 22 |  |
|  | Bilateral | 7 |  |
| Epileptogenic zone localisation, n | Temporal | 26 |  |
|  | Frontal | 11 |  |
|  | Insulo-opercular | 3 |  |
|  | Central-premotor | 3 |  |
|  | Posterior | 4 |  |
|  | Multilobar | 4 |  |
| Number of nodes, mean ± SD [median, min-max] | EZN | 6.36 ± 2.9 [5.5, 3-13] | |
| | PZN | 9.13 ± 3.6 [9, 3-13] | |
| | EEN | 13.88 ± 6.4 [14, 3-30] | |
| Surgical outcome (Engel class), n | I | 15 |  |
|  | II | 6 |  |
|  | III | 2 |  |
|  | IV | 3 |  |

Patients were categorised into two classes based on the anatomofunctional organisation of their epileptogenic zone: TLE (n = 26) where the EZN is mainly confined to temporal regions (including various types of TLE networks^70^ i.e. temporal lobe ‘plus’ classifications^71^), and NTLE (n = 25) where the EZN comprises either frontal, parietal or occipital regions (see Fig. 2A). Specific NTLE subclasses are noted in Table 1. Surgical outcome following 12-month follow-up was available for approximately half of all patients (n_TLE_ = 14, n_NTLE_ = 12) and used to derive seizure-free (Engel class I) and not seizure-free (Engel class II–IV) status. Over two-thirds of TLE patients with available data were deemed seizure-free (n = 10, 71.42%) compared to less than half of all patients with NTLE (n = 5, 41.67%). No statistically significant age differences between t-test groups were detected and no relationships to patient age presented in our results.

**Fig. 2:**
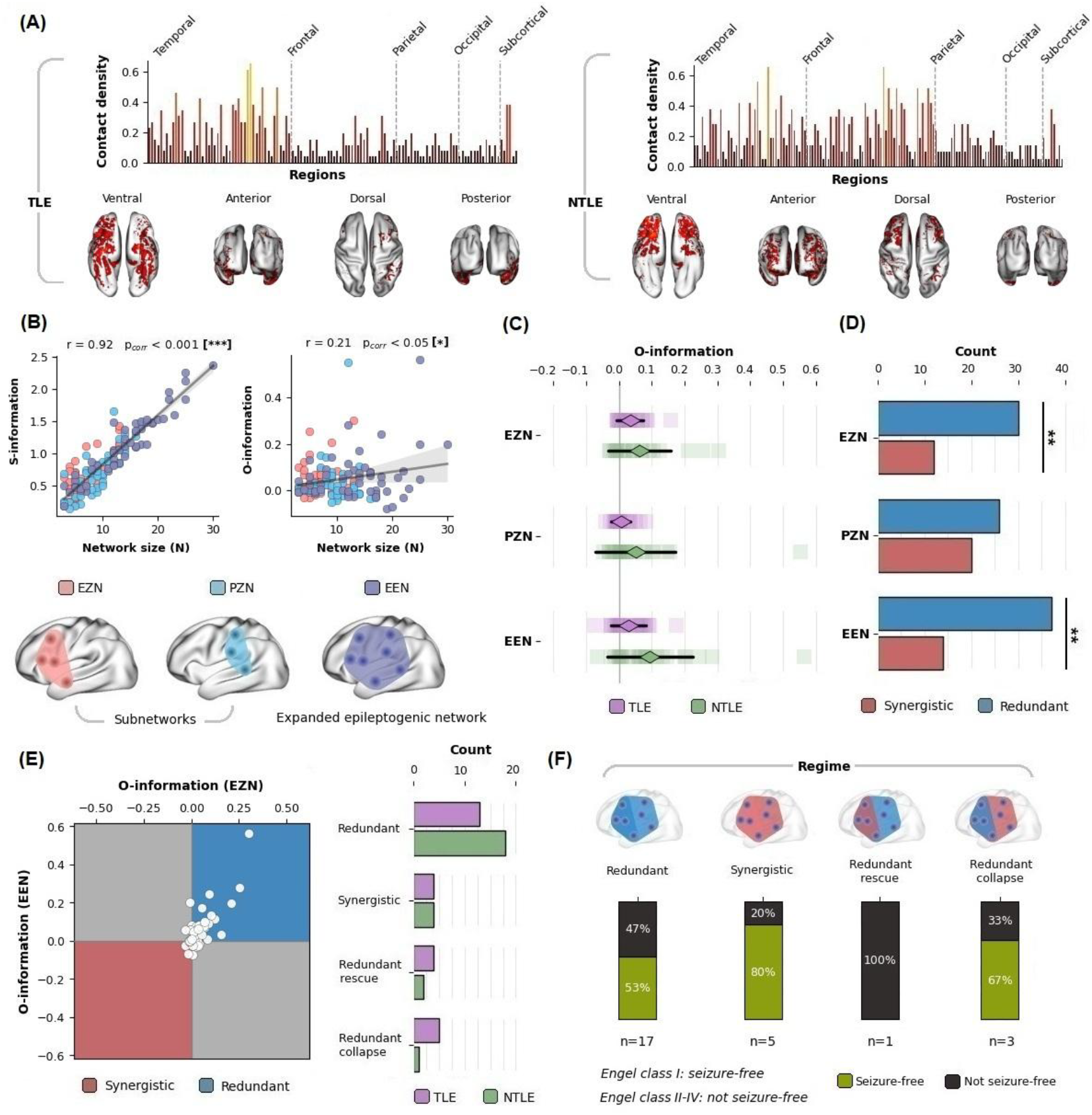
Higher-order information characterises temporal and extratemporal epileptogenic networks. **(A)** Bar charts comparing mean electrode contact density (normalised by group size) across brain regions, organised by lobe and projected onto surface plots for TLE and NTLE patients. Colour intensity represents group-level sampling density. **(B)** Dummy network partition visualisation and scatter plots displaying the normalised S- and O-information against network size (number of nodes *N*) with fitted regression lines and bootstrapped confidence intervals (1000 iterations; shaded grey band). **(C)** Distribution plot displaying individual network points (squares) with the mean (diamonds) and ± 1 SD (black lines) of the O-information per partition for each patient group. **(D)** Count plot comparing synergy-vs redundancy-dominated classifications between partitions, with binomial tests assessing deviations from a 1:1 split per network. **(E)** Symmetrical scatter plot of individual patients in O-information space defined by the EZN (x-axis) vs EEN (y-axis) and used to derive information-sharing regimes (synergistic [red], redundant [blue], redundant rescue and redundant collapse [both grey]) alongside a stacked count plot showing patient distributions stratified by focal epilepsy class. **(F)** Relative percentiles of seizure-free and not seizure-free patients following surgery for each information-sharing regime where available as represented by inset dummy networks. Asterix annotations indicate statistical significance (* p_corr_ < 0.05, ** p_corr_ < 0.01, *** p_corr_ < 0.001).

### Higher-order information

To understand how higher-order information varies with system size, we first examined to what extent the number of nodes in each epileptogenic network influences normalised S- and O-information. As seen in Fig. 2B, the S-information demonstrated a strong linear correlation with the number of nodes in the network (r = 0.92, p_corr_ < 0.001). In contrast, the O-information showed a weaker relationship to network size (r = 0.21, p_corr_ < 0.05). Given the size-dependent scaling of the S-information, we employed the signed O-information for comparisons between epileptogenic network partitions. As seen in Fig. 2C, a significant subject-level difference in O-information between each partition was detected following a Freidman test (Q = 15.21, p < 0.001). No significant group differences between partitions were detected, where mean O-information at the EZN (μ = 0.05 ± 0.07) and EEN (μ = 0.06 ± 0.1) was more positive compared to the PZN (μ = 0.03 ± 0.09) on average. Comparing focal epilepsy classes, overall, patients with NTLE displayed more positive O-information compared to patients with TLE (μ_TLE_ = 0.02 ± 0.04, μ_NTLE_ = 0.07 ± 0.12, t = -3.1, dof = 84.4, p = 0.002). No significant differences were detected between patients with TLE and NTLE at any network partition.

We performed binomial tests to clarify which type of information-sharing dominates each epileptogenic network partition during the interictal resting-state (H_0_: equal proportions of synergy-dominated and redundancy-dominated network).

We found that the EZN and EEN tended to be dominated by redundant information (see Fig. 2D), with significant differences in the proportion of synergy-dominated and redundancy-dominated networks per partition (% RED_EZN_ = 71.43, p_corr_ = 0.01; % RED_EEN_ = 72.55, p_corr_ < 0.01). No difference in proportions was detected at the PZN (% RED_EZN_ = 56.52). We employed Chi-squared tests of independence to determine test-retest stability between the two rs-fMRI acquisitions (see Supplementary Fig. 3). We found that the EZN and EEN demonstrated stable proportions (χ^2^_EZN_ = 0.06, p = 0.81; χ^2^_EEN_ = 0.05, p = 0.83) whilst the PZN showed greater variability between acquisitions (χ^2^_PZN_ = 5.43, p = 0.02, p_corr_ = 0.06) indicated by a higher proportion of synergy-dominated networks in the first rs-fMRI run (% RED_PZN_ = 45.65).

We assigned patients to regimes based on O-information of the EEN and (where possible) the EZN to characterise how information-sharing may differ between regions of seizure onset and the wider pathological network. Each patient was classified under one of four possible combinations (see Fig. 2E): redundant (Ω_EZN_ > 0, Ω_EEN_ > 0), synergistic (Ω_EZN_ < 0, Ω_EEN_ < 0), redundant collapse (Ω_EZN_ > 0, Ω_EEN_ < 0) or redundant rescue (Ω_EZN_ < 0, Ω_EEN_ > 0). Redundant collapse describes a shift from predominantly redundant information within the EZN to predominantly synergistic information when the PZN is included, whereas redundant rescue describes the opposite transition. Plotting the O-information of the EEN against that of the corresponding EZN, deviations from zero were generally skewed towards positive values along both axes (see Fig. 2D). Most patients exhibited a redundant regime (n = 31, 60.78%; n_TLE_ = 13, n_NTLE_ = 18 [7 frontal, 3 central-premotor, 3 insulo-opercular, 3 posterior, 2 multilobar]) whilst a minority were synergistic (n = 8, 15.69%; n_TLE_ = 4, n_NTLE_ = 4 [2 frontal, 1 posterior, 1 multilobar]. The remaining patients were divided between redundant rescue (n_TLE_ = 4, n_NTLE_ = 2 [all frontal]) and redundant collapse (n_TLE_ = 5, n_NTLE_ = 1 [multilobar]). No significant association was observed between focal epilepsy class and information-sharing regimes following a Chi-squared test of independence. Relative percentiles of seizure free and not seizure-free patients per information-sharing regime and focal epilepsy class are also reported in Fig. 2F and Fig. 4E respectively.

### Relationship to conventional connectivity metrics

To explore how information-sharing is related to a network’s connectivity profile and varies across partitions, we computed seven conventional metrics and compared redundancy and synergy-dominated epileptogenic networks. Across all network-level connectivity metrics (i.e. the average between all region pairs in a network for each metric), the EZN, PZN and EEN demonstrated markedly distinct connectivity profiles following Freidman tests (all p_corr_ < 0.001). Subsequent univariate testing did not reveal any statistically significant differences between synergy- and redundancy-dominated networks across any partition following corrections for multiple comparisons. Nevertheless, uncorrected p-values indicated observable differences in mean streamline weight, mean FC and mean centroid Euclidean distance particularly at the EZN (see Fig. 3A).

**Fig. 3:**
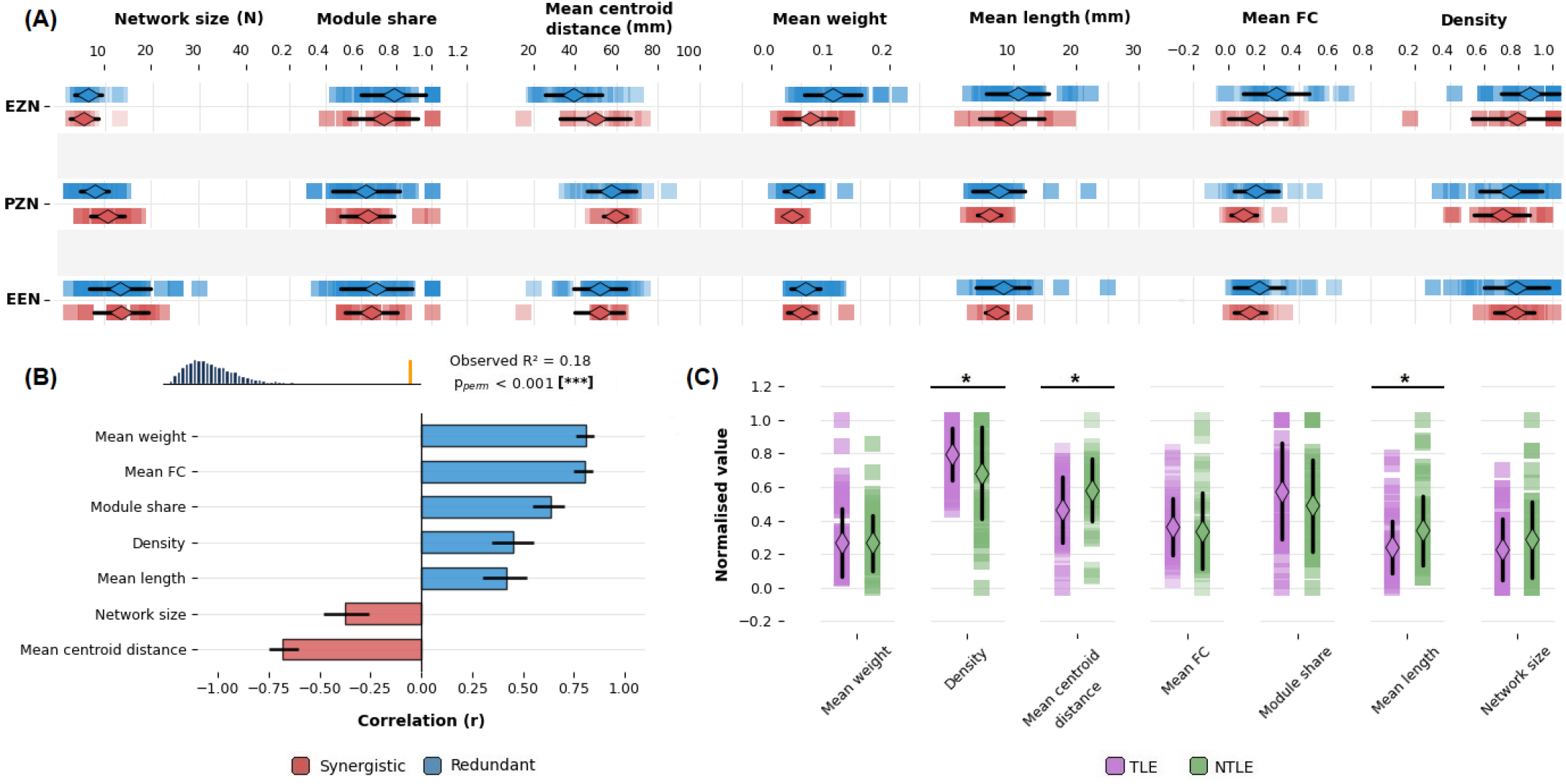
Synergy- and redundancy-dominated epileptogenic networks display distinct connectivity profiles. **(A)** Seven-panel grid showing structural and functional connectivity metrics across epileptogenic network partitions stratified by dominance category (redundant: blue, synergistic: red). Each panel shows individual data points (squares) overlaid with mean ± 1 SD (black lines) and the deviation mean (diamonds). **(B)** Bar plot displaying Pearson correlations between each standardised connectivity metric and PLS component-1 scores coloured by direction (blue [r > 0, associated with redundancy] or red [r < 0, associated with synergy]). Error bars represent 95% confidence intervals of r derived from Fisher z-transforms. Inset histogram contains permuted R^2^ values (grey bars) with a single vertical line (orange) marking the observed R^2^. **(C)** Distribution plot displaying normalised connectivity metrics (min-max scaling: 0-1) stratified by focal epilepsy class across all seven connectivity measures. Asterix annotations indicate statistical significance for group differences in non-normalised connectivity metrics (* p_corr_ < 0.05, ** p_corr_ < 0.01, *** p_corr_ < 0.001).

Moving to multivariate testing, we employed PLS regression with O-information as the continuous outcome variable and the seven network connectivity metrics as standardised predictors to identify which features collectively indicated synergy or redundancy dominance. Observing all available networks (n = 113), a single latent component was selected via five-fold cross-validation (R^2^ = 0.06) and demonstrated significant predictive power following permutation testing (R^2^ = 0.18, p_perm_ = 0.0002). As seen in Fig. 3B, PLS component loadings revealed redundancy-associated features contained the strongest positive contributions. These included mean weight (r = 0.81) and mean FC (r = 0.8) followed by module share (r = 0.63), density (r = 0.45), and mean length (r = 0.42). In contrast, mean centroid distance (r = -0.68) and network size (r = -0.38) displayed negative loadings, confirming an association with synergy-dominated epileptogenic networks. This process was repeated using separate loadings from EZN, PZN and EEN samples respectively to visualise the relative contributions of each partition to our combined PLS result (see Supplementary Fig. 4).

Finally, to clarify global connectivity differences between TLE and NTLE networks in our dataset, we pooled all epileptogenic networks and compared differences in connectivity metrics between the two focal epilepsy classes. We found that NTLE networks demonstrated more dispersed spatial arrangements than TLE networks (see Fig. 3C), with significant differences in mean centroid Euclidean distance (μ_TLE_ = 47.44 mm ± 13.7, μ_NTLE_ = 55.69 mm ± 13.17, t = -3.24, dof = 108.4, p_corr_ = 0.01) and mean streamline length (μ_TLE_ = 7.35 mm ± 3.57, μ_NTLE_ = 9.67 mm ± 4.86, t = -2.84, dof = 89.8, p_corr_ = 0.02). On the other hand, TLE networks displayed significantly elevated connection density on average (μ_TLE_ = 0.83 ± 0.13, μ_NTLE_ = 0.74 ± 0.23, t = 2.53, dof = 75.6, p_corr_ = 0.03). No significant differences between the two groups were observed across any other metric.

#### Dual Z-score comparisons

To understand the extent to which the higher-order information of epileptogenic networks could be pathology-related, we derived Z-scores from two distinct reference distributions and compared focal epilepsy classes (see Fig. 4A). For Z_w_-scores, a single patient network failed to yield 1000 null samples and was discarded from further analysis. Upon histogram inspection, Z_w_-score distributions were remarkably Gaussian whereas Z_b_-score distributions exhibited more variability (see Supplementary Fig. 5). Assuming normality across both distributions, significant deviations from the mean of each distribution were considered at |*z*| > 1.96.

**Fig. 4:**
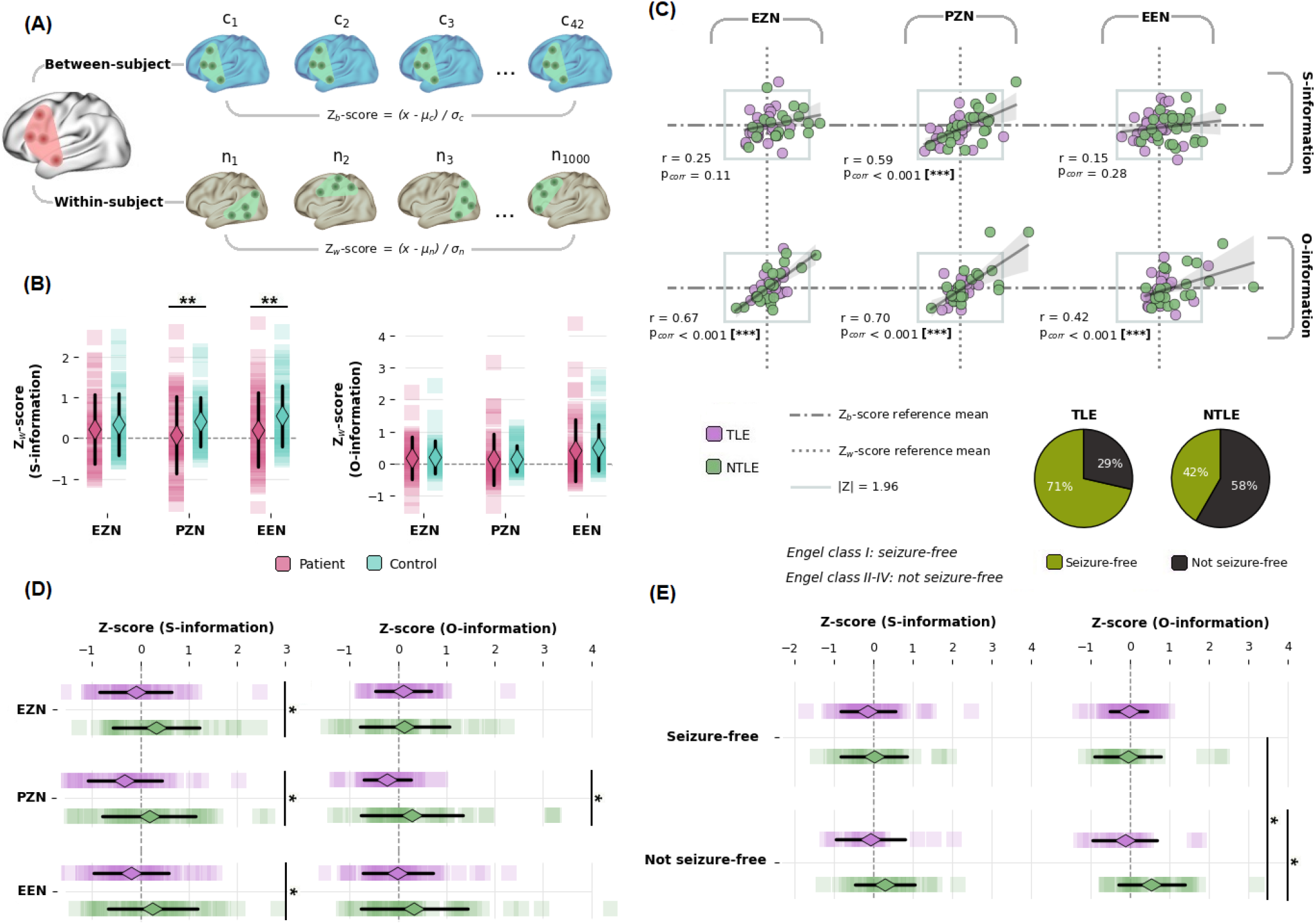
Combined between- and within-subject comparisons uncover differences in information-sharing between temporal and extratemporal networks and surgical outcomes. **(A)** Overview of Z_b_- and Z_w_-score calculation from homologous healthy control (c) and null NI/NE (n) network distributions represented by dummy networks **(B)** Distribution plot comparing S- and O-information Z_w_-scores between patients and controls across the three epileptogenic network partitions. The grey dash-dot line indicates the null average, while diamonds and black lines represent the mean and ± 1 SD respectively. Paired t-tests (matched subject network pairs only; average number of computed pairs reported) compare patient vs control mean Z_w_-scores per metric and partition. **(C)** Correlation analysis plotting Z_b_-(y-axis) against Z_w_-scores (x-axis). Each point represents a single network coloured by focal epilepsy class. Grey reference encasings at ±1.96 SD mark deviation significance bounds. Linear regression lines are fitted to each network scatter and 95% confidence intervals computed via bootstrap (1000 iterations; shaded grey band). Reference grey dash-dot and dotted lines at zero reflect the healthy control and null mean for Z_b_- and Z_w_-score calculation respectively. **(D)** Combined Z_b_- and Z_w_-scores for each metric across each partition, with the dashed grey reference lines indicating deviations of zero. **(E)** Distribution plot displaying combined Z-scores for patients who underwent surgery following intracranial recordings. S- and O-information deviations are plotted separately (along with their respective means and ± 1 SD) stratified by focal epilepsy class and split by surgical outcome (seizure-free vs not seizure-free). Attached pie charts indicate the relative percentages of seizure-free and not seizure-free patients in our TLE and NTLE cohorts. Asterisk annotations indicate statistical significance following paired or unpaired t-testing (* p_corr_ < 0.05, ** p_corr_ < 0.01, *** p_corr_ < 0.001)

First, we compared patient Z_w_-scores against matched homologous control network Z_w_-scores. The mean number of successfully computed homologous control network Z_w_-scores per patient prior to averaging (out of a possible 84 [42 controls × 2 runs]) were as follows: μ_EZN_ = 53.57 ± 19.1, μ_PZN_ = 72.67 ± 16.11 and μ_EEN_ = 60.41 ± 19.18. Applying paired t-tests, patients demonstrated significantly reduced S-information Z_w_-scores relative to controls at the PZN (μ_patient_ = 0.08 ± 0.95, μ_control_ = 0.4 ± 0.61, t = -3.12, dof = 45, p_corr_ = 0.01) and EEN (μ_patient_ = 0.2 ± 0.91, μ_control_ = 0.54 ± 0.74, t = -4.01, dof = 50, p_corr_ = 0.001) with no statistically significant difference detected at the EZN. No significant patient-control differences emerged for the O-information Z_w_-scores following the same matched design (see Fig. 4B)

Comparing Z_b_- and Z_w_-scores directly, as shown in Fig. 4C, O-information deviations showed strong positive correspondence across all partitions (r_EZN_ = 0.67, r_PZN_ = 0.70, r_EEN_ = 0.42, all p_corr_ < 0.001). S-information deviations demonstrated a similar relationship restricted to the PZN (r = 0.59, p_corr_ < 0.001) with no significant correspondence detected at the EZN or EEN. The details of individual patients who exhibited significant Z_b_- or Z_w_-scores are highlighted in Supplementary Table 2.

Pooling all Z_b_- and Z_w_-scores, as shown in Fig. 4D, patients with NTLE displayed significantly more positive S-information deviations than patients with TLE across all partitions (EZN: t = -2.33, dof = 76, p_corr_ = 0.03; PZN: t = -2.74, dof = 90, p_corr_ = 0.02; EEN: t = -2.61, dof = 96, p_corr_ = 0.02). For the O-information, deviations amongst NTLE networks were more positive on average (μ_TLE_ = -0.03 ± 0.62, μ_NTLE_ = 0.26 ± 1.02, t = -2.89, dof = 220.6, p_corr_ = 0.004) and differed significantly from TLE at the PZN only (μ_TLE_ = -0.22 ± 0.49, μ_NTLE_ = 0.29 ± 1.04, t = -3.01, dof = 90, p_corr_ = 0.02). Correlation analyses revealed no significant association between S-and O-information deviations and network size (see Supplementary Fig. 6).

Finally, stratifying by surgical outcome amongst available patients (n = 26), not seizure-free patients with NTLE exhibited significantly more positive O-information deviations than seizure-free patients with NTLE (μ_SF_ = -0.05 ± 0.85, μ_NSF_ = 0.54 ± 0.84, t = -2.83, dof = 62, p_corr_ = 0.03), an effect that was absent amongst TLE cases (see Fig. 4E). The same elevation distinguished NTLE from TLE cases specifically within the not seizure-free group (μ_TLE_ = -0.13 ± 0.82, μ_NTLE_ = 0.54 ± 0.84, t = -2.76, dof = 50, p_corr_ = 0.03) whilst no difference between focal epilepsy classes was found amongst seizure-free patients. No significant differences in S-information deviations were observed across surgical outcome or focal epilepsy class.

## Discussion

DRFE is understood as a complex epileptogenic network, comprising groups of interacting brain regions between which information is shared both during and between seizure events. Capturing global dynamics, multivariate information theory reveals higher-order interactions emerging from epileptogenic networks in presurgical rs-fMRI. Our application of the S- and O-information offers novel descriptions of epileptogenic network organisation that show a relationship to conventional MRI-based connectivity measures and may prove useful to the epileptologist during presurgical planning.

### The redundant epileptogenic network

The O-information defines the balance between synergistic and redundant information-sharing in a system. Confined mainly to the EZN and EEN, most patients displayed networks dominated by redundancy (see Fig. 2D). These networks exhibited stronger structural and functional connections between regions compared to those dominated by synergy (see Fig. 3B), in line with recent work^33,34,72^. Notably, this effect was prevalent at the epileptogenic zone, where regions may be neighbouring as in TLE, or interconnected by longer-range anatomical projections as in NTLE. Redundant information-sharing when epileptogenic regions are contained in a system could be related to the underlying anatomofunctional properties of seizure networks. Previous work has highlighted maladaptive or abnormally elevated connectivity between regions of seizure onset^73,74^. Indeed, a similar retrospective MRI investigation found structural connectivity to be preserved both within and between EZN and PZN regions when compared to healthy controls^75^. Seizures themselves are theorised to progressively align with WM connections^76^ which serve as individualised scaffolds for modelling ictogenesis^3,77^. Coupled with findings from electrophysiology describing redundancy and maintained FC at the epileptogenic zone^78-80^, redundant information-sharing during the interictal resting-state offers a tangible reflection of epileptogenic network organisation. Investigating how the O-information evolves through the ictal process will be important in determining whether redundancy dominance reflects a stable network property or a dynamic feature of seizures moving forward.

### Synergies and network complexity in the epileptic brain

Our results demonstrate how epileptogenic networks can also be dominated by synergistic information-sharing. As suggested in recent work, synergy dominance was predicted by the presence of dispersed or spatially isolated regions in larger networks^71^. Specifically, cases of a synergy-dominated EZN appeared restricted to frontal NTLE and likely involved components of the default mode network, a known area of synergistic information in the brain^30,33,81^. Understanding how information-sharing in epilepsy relates to canonical functional properties could serve to decouple abnormal network effects from those typical of the resting-state. In addition, a noted hallmark of synergy-dominated networks is the involvement of intermodular brain regions; that is, those nodes or hubs exhibiting high centrality attributes^33,72^. Hub dysfunction is well described in both focal and generalised epilepsies^82,83^ and has expanded to include subcortical malformations^44,84,85^. Whilst elucidating the involvement of specific regions is beyond the scope of the current study, future work may wish to examine the contribution of hubs to the information-sharing properties of epileptic brain networks.

We also explored how the overall strength of interdependencies, the S-information, differs between networks of epileptic and non-epileptic regions. The S-information is closely linked to the Tononi-Sporns-Edelman (TSE) complexity^30,32^, a measure which is maximised during moments of global integration and local segregation in a system^29,86,87^. Put differently, the complexity of a network is low when its nodes show no coherency between them or, oppositely, when those same nodes are too constrained in their collective activity. Our Z-score analysis revealed differences between the S-information Z^w^-scores of epileptogenic and homologue-matched networks, with the PZN and EEN demonstrating weaker information-sharing compared to the rest of the brain than the same regions in healthy controls (see Fig. 4B). Given the disease context, these networks may exhibit an impaired capacity for the enriched, complex interactions characterising healthy brains^88,89^. Interestingly, no differences in the O-information were observed following the same analysis, suggesting that information-sharing type is relatively preserved whilst the strength of interdependencies surrounding that information is diminished at the PZN and EEN. Future research may wish to ask whether the S-information is linked to certain ictal signs and comorbidities such as alterations in awareness^90^ in epileptic brain networks.

### Clinical value of multivariate information theory in presurgical resting-state fMRI

We attained deviation scores from two reference distributions to understand the degree to which higher-order information could be pathology-related in epileptogenic networks. Resulting Z_b_- and Z_w_-scores, particularly in the case of the O-information, showed remarkable correspondence across all partitions (see Fig. 4C). Such agreement between two independently constructed distributions suggests that our population-based and idiosyncratic models capture consistent shifts in information-sharing that are linked, at least in part, to the DRFE disease context.

Our combined comparative framework identified at least one epileptogenic network where the S- or O-information deviated significantly from the reference distribution in about 20% of patients (see Supplementary Table 2). These Z-scores were most concentrated at the EZN and EEN and comprised mainly patients presenting with NTLE. Almost all detected patients displayed a redundant information-sharing regime and two of a cohort total of five patients with a surgical outcome of Engel class III–IV were also detected. In addition, our sensitivity analysis revealed that widening of the mean FC acceptance range during null distribution construction led to an increased number of significant deviations (see Supplementary Fig. 2C). Controlling for bivariate connectivity, epileptogenic networks may thereby exhibit pathological higher-order properties that are not easily decomposed to pairwise components. Aligning with recent work charting the added value of higher-order interactions to BOLD time series analysis^91^, beyond pairwise exploration of rs-fMRI data could offer a complementary network-based approach in personalised epileptology^92,93^. Considering the associations between FC and redundant information^34,91^, future benchmarking will prove pivotal in clarifying advantages of higher-order frameworks over conventional methods in DRFE.

Considering correlations with the normalised S- and O-information and the number of nodes *N*, we combined Z-score pairs to examine differences in information-sharing independent of network size. We found differences between S-information deviations, with NTLE networks showing stronger interdependencies between regions relative to what either their own brain or healthy controls would predict, whereas TLE networks were mostly below expected strengths (see Fig. 4D). Dovetailing our connectivity findings (see Fig. 3C), NTLE networks appear to be not only larger and more spatially distributed than TLE networks, but also more functionally complex. O-information deviation differences, conversely, did not present at the EZN or EEN, suggesting that the synergy-redundancy balance along propagation pathways is where TLE and NTLE tend to diverge. Together, these findings support the view that NTLE encompasses a more diverse interictal resting-state profile than TLE, the likes of which may be captured by measures of multivariate information.

Our retrospective MRI analysis occupies a critical stage of the DRFE clinical pathway between intracranial recordings and potential surgery. Pooling all Z-score pairs revealed the clearest outcome-related finding: patients with NTLE who were not seizure-free following surgery displayed more positive O-information deviations than seizure-free patients, with the same elevation also distinguishing them from not seizure-free TLE cases (see Fig. 4E). Pointing to an association between elevated redundancy and postoperative seizure freedom, the sharing of copied information between epileptogenic network regions may contribute to a robust functional architecture that is comparatively resistant to localised disruptions^35,94,95^. Under this interpretation, resection or ablation of individual EZN regions may be less effective in NTLE, where redundancy in the expanded network could sustain pathological activity (see Fig. 2F), possibly by way of abnormal structural connections.^96^ In contrast, O-information deviations in seizure-free NTLE were considerably closer to those in TLE, consistent with a synergy-redundancy balance that could be more amenable to network disconnection. Given our group sizes, these observations should be regarded as preliminary; though they nevertheless highlight how multivariate information theory, and redundancy measures in particular, can offer candidate biomarkers of network organisation and surgical outcome in DRFE.

### Methodological choices and limitations

This study contains several methodological choices and limitations to be clarified. First, our rs-fMRI analysis relies on the gold-standard clinical definition of epileptogenic and propagator zones from SEEG. With electrode implantation hypothesis-driven and spatially limited, inevitable biases are introduced whereby pathological tissue outside sampled regions may remain unexplored. Likewise, regions assigned to the EZN were considered entirely epileptogenic for network construction and null model sampling despite epileptogenic tissue possibly occupying only a small component of each contacted region. Future work combining SEEG findings with voxel-wise approaches could provide more precise definitions of EZN and PZN regions in retrospective MRI analysis.

We employed a simple null model using BOLD time series data and region centroid coordinates to compare epileptogenic networks to subsets of NI/NE regions of equivalent size, similar spatial dispersion and comparable mean FC. Whilst this approach controls major spatiotemporal confounds and maximises the sampling space, it does not account for structural topology, nodal properties or resting-state network membership, and is limited to VEP atlas regions. We were also unable to compute null distributions for a single patient and several healthy controls due to the constraints and size requirements of our model, which likely introduced some bias into our paired Z_w_-score comparisons. Future null models incorporating structural connectivity and different parcellation schemes could provide more biologically and computationally stable reference distributions.

We focused on the network-level S- and O-information to obtain global descriptions of information-sharing whilst avoiding the combinatorial explosion associated with lower-order multiplets. This strategy enabled tractable patient-specific null modelling but sacrificed local interactions between subsets of regions, edgewise effects and node-level contributions. Our analysis was also confined to static interictal resting-state data averaged across acquisitions rather than dynamical higher-order interactions. With the synergy-redundancy balance shown to fluctuate at rest^34,97^, future work could employ time-resolved approaches to reveal network states that may be obscured by summary measures.

Finally, our clinical analysis should be interpreted cautiously due to its interictal setting and modest sample size. Prospective studies of seizure recordings that combine measures of multivariate information with larger multicentre cohorts will be important in establishing the robustness and clinical utility of these methods moving forward.

## Data availability

Data has not been published due to sensitive information that could compromise patient confidentiality. Anonymised data may be made available from the corresponding author upon reasonable request.

## Code availability

VEP parcellation and time series analysis code for the current study is openly available at https://github.com/jacob-theknight/smk-epilepsy-ts. Code development was assisted by GitHub Copilot Pro in Visual Studio Code (v1.127.0) for code suggestions and debugging. The implementations of the S-information and O-information are based on the open-source Python toolbox HOI available at https://brainets.github.io/hoi/.

## Author contributions

J.K conceived the study idea and carried out the formal analysis with contributions from M.N and R.H. MRI data was preprocessed by H.D and J.K. SEEG implantations and recordings were performed by medical doctors J.M and F.B. EZN, PZN and NI electrode contacts were determined by J.M and F.B, and CT-MRI co-registrations were performed by S.M. M.N, J.R, A.B, M.G and R.H contributed to the final study design and interpretation. J.K wrote the original draft with valuable revisions from R.H. All authors contributed to the final draft review and editing.

## Acknowledgments

We would like to thank Patrick Viout, Lauriane Pini, Claire Costes, and Veronique Gimenez for MRI data acquisition and study logistics. We would also like to extend our thanks to Enrico Amico for his valuable review and advice regarding the manuscript. We are grateful to all the patient and control participants who agreed to take part in this study.

## Funding sources

This study received funding from the French government under the Programme Investissements d’Avenir and France 2030 (ANR-16-CONV000X and ANR-17-EURE-0029), Excellence Initiative of Aix-Marseille University A*MIDEX (AMX-19IET-004), 7TEAMS Chair, EPINOV (ANR654 17-RHUS-0004) and the European Union’s Horizon 2020 Framework Program (785907 and 945539). This work was performed by members of France Life Imaging network (ANR-11-INBS-0006), Marseille Imaging Institute (AMX-19IET-002) and NeuroMarseille Institute (AMX-19IET-004). Author J.K is supported by the Agence Nationale de la Recherche (ANR-24659 CPJ1-0143-01).

## Competing interests

The authors declare no competing interests.

## Supplementary material

**Supplementary Table 1:** Summary of terminology and corresponding abbreviations used in reporting group comparisons.

| Category | Terminology | Abbreviation |
| --- | --- | --- |
| Epileptogenic network partition | Epileptogenic zone network | EZN |
|  | Propagator zone network | PZN |
|  | Expanded epileptogenic network | EEN |
| Focal epilepsy type | Temporal lobe epilepsy | TLE |
|  | Non-temporal lobe epilepsy | NTLE |
| Information-sharing type | Synergy-dominated | SYN |
|  | Redundancy-dominated | RED |
| Surgical outcome | Seizure-free | SF |
|  | Not seizure-free | NSF |

**Supplementary Table 2:** Details of patients who exhibited at least one significant Z_b_- or Z_w_-score across any epileptogenic network or metric.

| Subject | Identified network(s) | Identified metric(s) | Deviation direction | Age | Sex | Focal epilepsy subtype | Laterality | MRI-negative | Surgical outcome | Identified regime |
| --- | --- | --- | --- | --- | --- | --- | --- | --- | --- | --- |
| 007 | EZN, EEN | $\Omega$ | + | 40 | M | Frontal | R | No | I | Redundant |
| 046 | EZN | $\Sigma$ | + | 40 | F | Central-premotor | R | No | IV | Redundant |
| 060 | EZN | $\Omega$ | + | 30 | M | Temporal | L | Yes | n.a. | Redundant |
| 070 | EZN | $\Sigma$ | + | 18 | F | Temporal | R | Yes | I | Redundant |
| 122 | EZN, EEN | $\Sigma, \Omega$ | + | 15 | M | Posterior | R | No | n.a. | Redundant |
| 097 | PZN | $\Sigma$ | + | 22 | M | Temporal | R | No | I | Rescue |
| 158 | PZN | $\Omega$ | + | 46 | F | Multilobar | L | No | III | Redundant |
| 155 | PZN, EEN | $\Sigma, \Omega$ | + | 42 | M | Central-premotor | L | Yes | n.a. | Redundant |
| 061 | EEN | $\Omega$ | + | 27 | F | Temporal | R | Yes | n.a. | Redundant |
| 083 | EEN | $\Sigma$ | - | 16 | M | Temporal | Bilateral | No | n.a. | Redundant |

**Supplementary Fig. 1:**
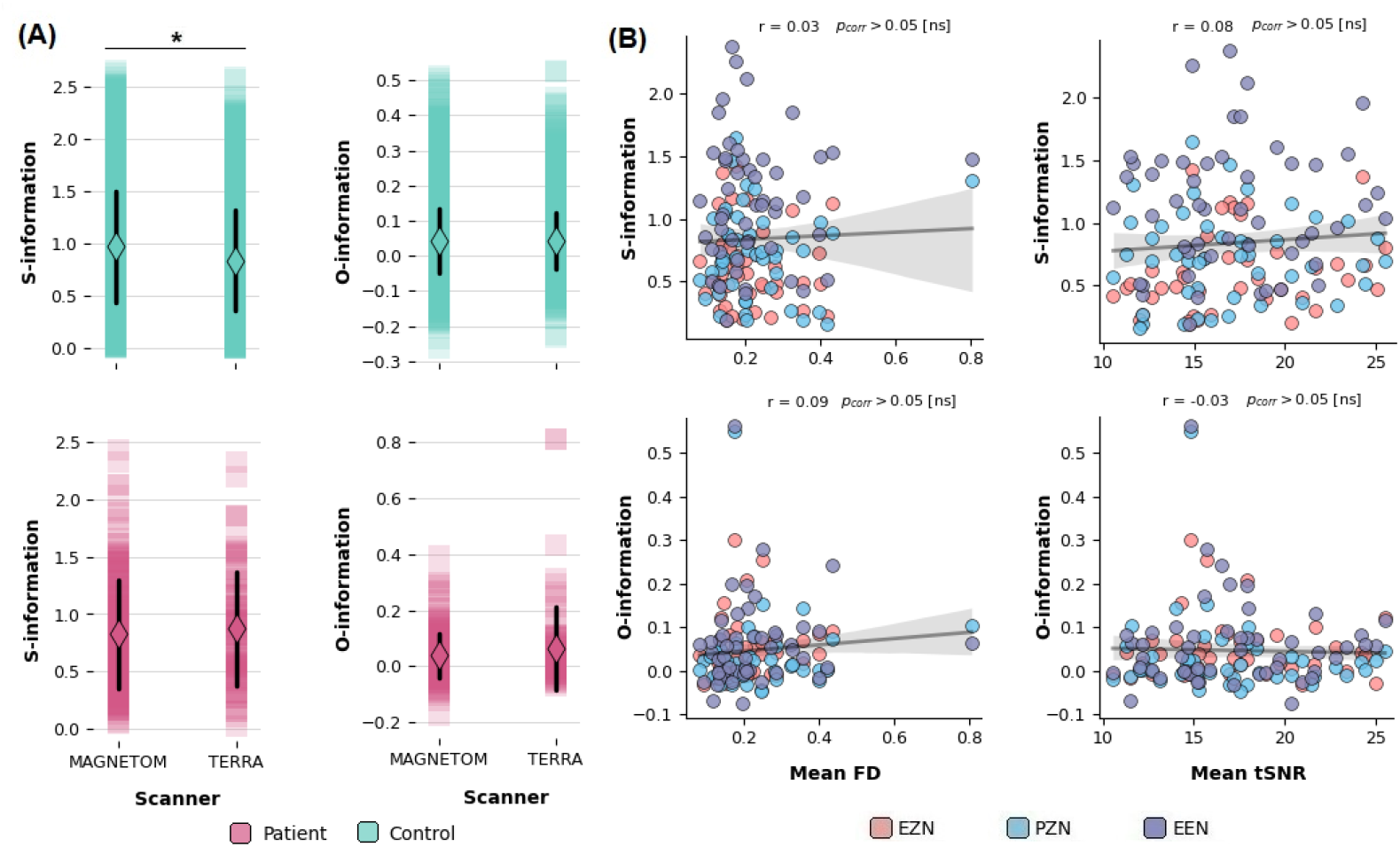
Post-hoc quality control assessing associations between the normalised S- and O-information and scanner differences alongside global IQMs. **(A)** Distribution plots of normalised measures of multivariate information stratified by the two systems used for rs-fMRI within our cohort (MAGNETOM vs TERRA). Diamonds and black lines indicate means and ± 1 SD respectively, whilst each data point (square) represents S- and O-information values derived from epileptogenic networks. A total of 60 subjects (37 patients, 23 controls) were scanned by our Magnetom system and a total of 33 subjects (14 patients, 19 controls) most recently by our Magnetom Terra system. Healthy control homologue networks scanned by the Magnetom system exhibited significantly greater S-information values than those scanned by the Magnetom Terra system (marked by asterix), confirmed by a one-way ANOVA (F = 97.24, dof = 1, p < 0.0001). The same effect failed to present amongst patient networks despite corresponding acquisition parameters and therefore did not likely bias our final Z_w_-score comparisons. **(B)** Scatter plots with regression lines showing the relationship between measures of multivariate information and whole-brain mean framewise displacement (FD) and mean temporal signal-to-noise ratio (tSNR). Linear regression with 95% confidence intervals estimated via 1000 bootstrap iterations of resampled data pairs. No significant association was detected between IQMs and the normalised S- and O-information, indicating that global functional image quality did not likely bias our results.

**Supplementary Fig. 2:**
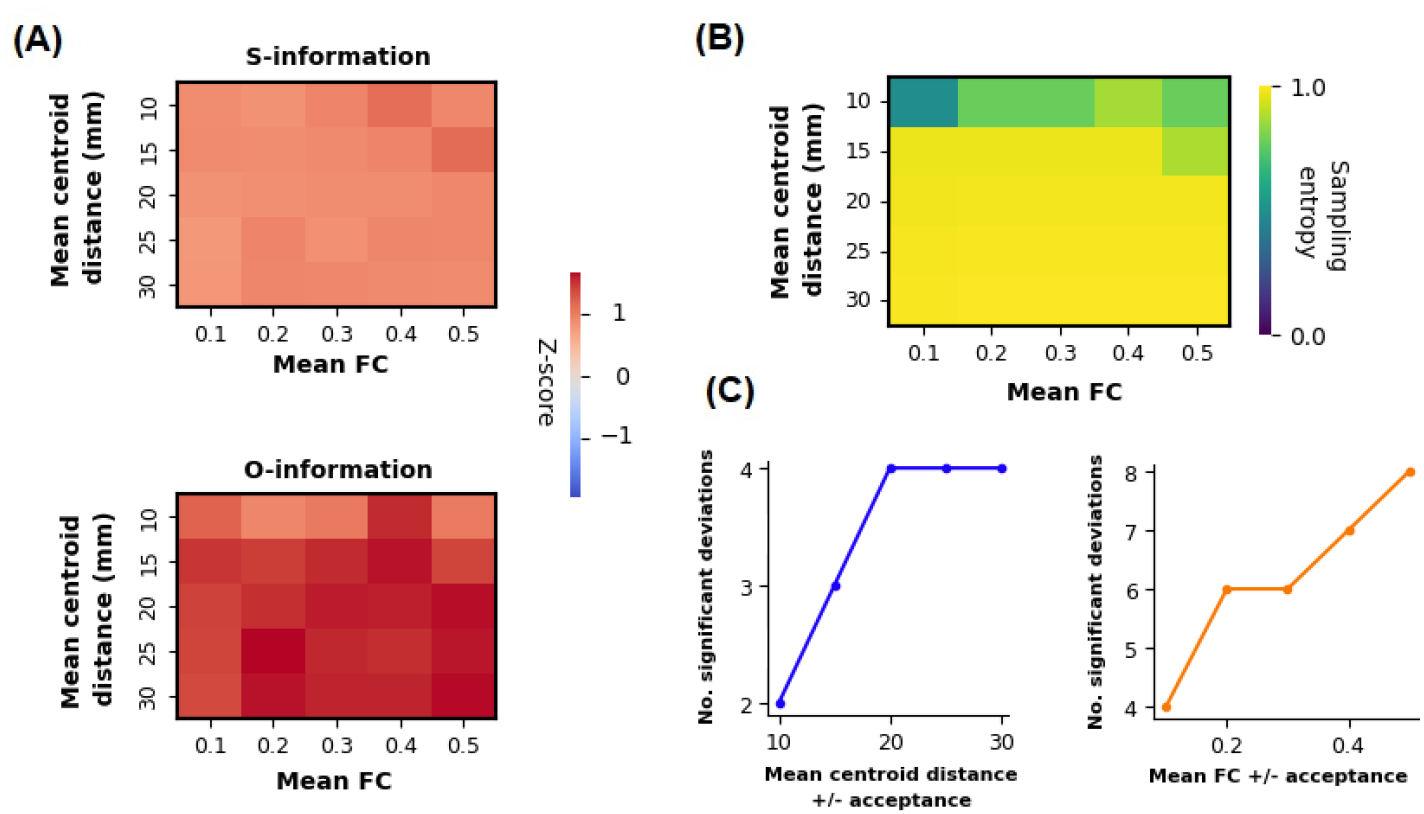
The S- and O-information Z_w_-scores are influenced by spatiotemporal null model constraints. **(A)** Two heatmaps showing mean Z_w_-scores for the two metrics across varying spatiotemporal matching ranges. Rows represent mean Euclidean centroid distance (10-30mm) and columns represent mean FC ± ranges for network acceptance (0.1-0.5) into the null distribution. Widened ranges tended to yield slightly elevated Z_w_-scores amongst a subset (n = 10) of patient EEN networks detected by our comparative framework. **(B)** Heatmap displaying mean sampling entropy values across the same null model parameter space. Sampling entropy was computed as the Shannon entropy of the frequency distribution of regions across accepted null network combinations, normalised by the logarithm of the number of unique regions. Widened ranges, as expected, led to augmented sampling entropy in our null model. **(C)** Line plots showing the number of significant deviations (|*z*| > 1.96) in S- and O-information as a function of the mean centroid distance (blue) and mean FC (orange) ± ranges across all patient EEN networks. Widened mean FC acceptance ranges appeared to increase null model sensitivity whereas mean centroid distance increases exhibited a weaker effect on deviation magnitude.

**Supplementary Fig. 3:**
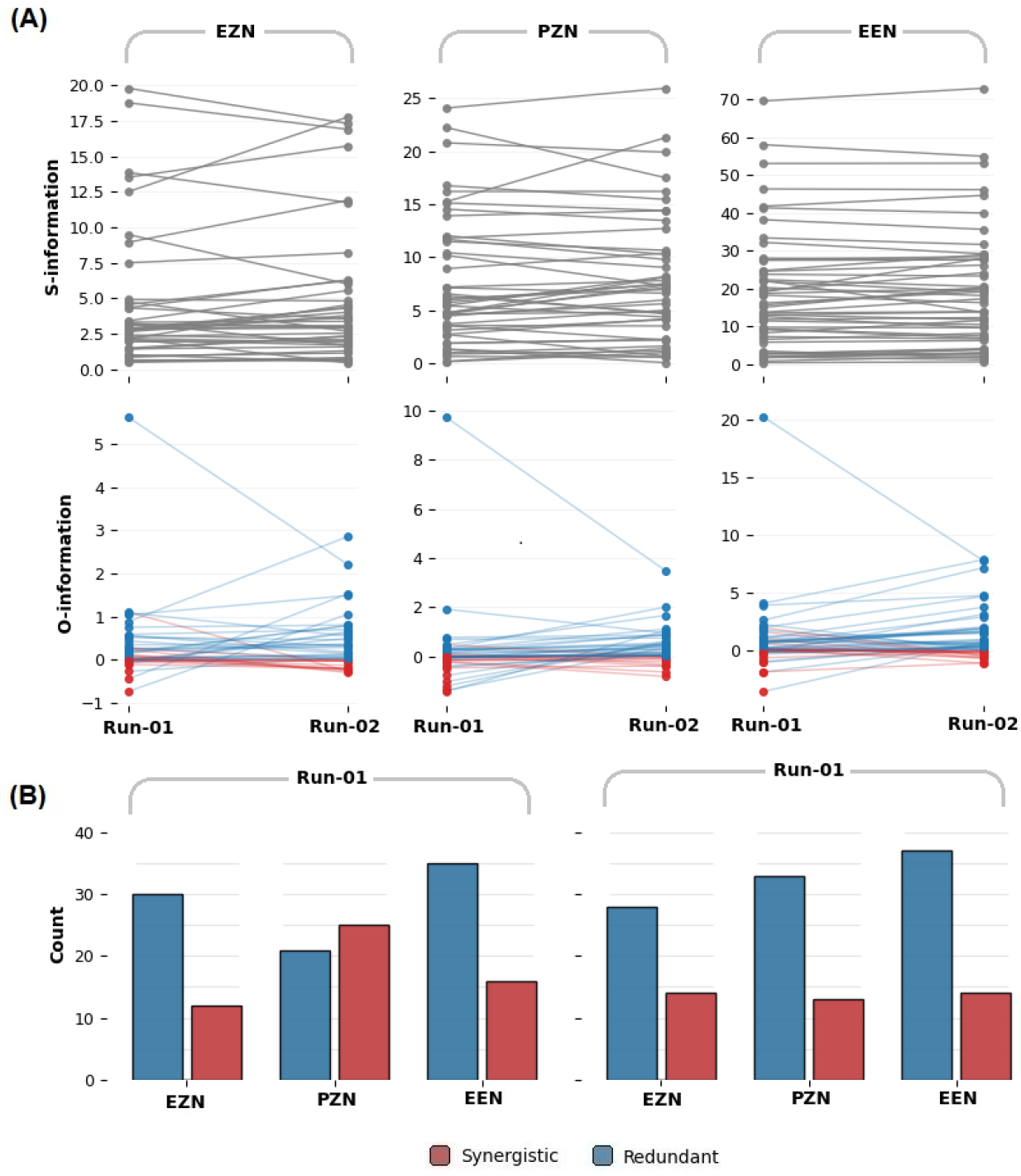
Synergy- and redundancy-dominated classifications display test-rest stability at the epileptogenic zone and expanded network. **(A)** Spaghetti plots displaying the raw S- and O-information of each epileptogenic network recorded over the two rs-fMRI acquisitions (Run-01 and Run-02) **(B)** Paired count plots for each acquisition showing the distribution of synergy-dominated vs redundancy-dominated networks derived from the raw O-information. Chi-square tests were used to assess run differences (χ^2^ values reported).

**Supplementary Fig. 4:**
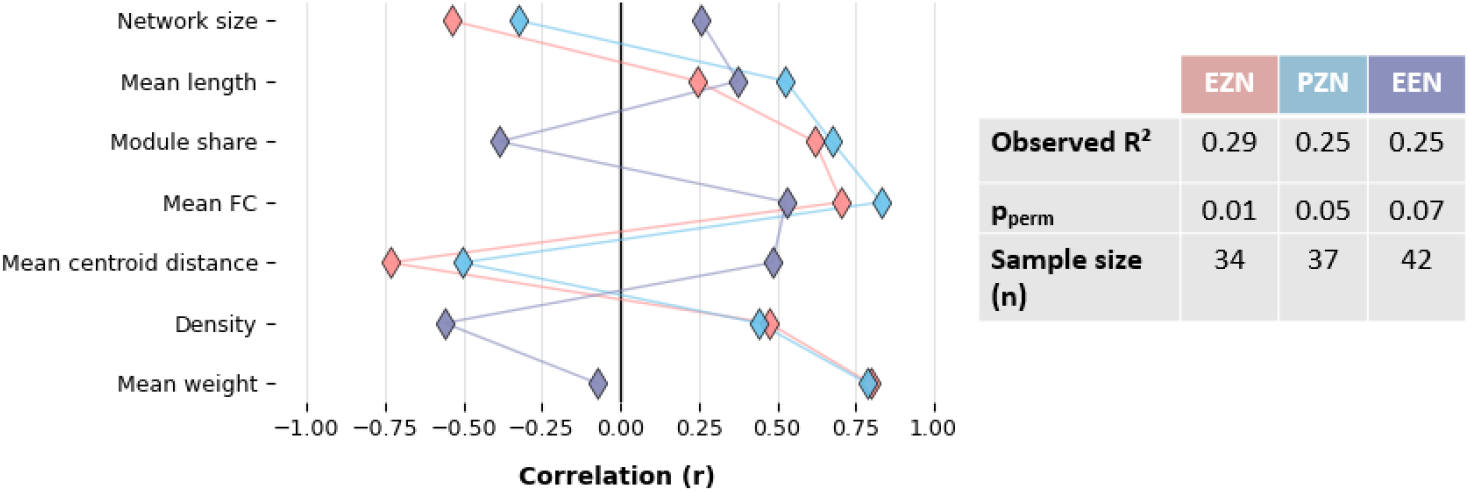
Individual PLS regression loadings for each connectivity metric reveals underlying epileptogenic network contributions. Each metric is represented by three diamond markers (one per epileptogenic network partition) with connector lines to visualise the component-1 loading score pattern. Positive and negative loadings indicate metrics associated with redundancy and synergy respectively. Individual PLS models were fitted independently for the EZN, PZN and EEN. Network-specific R^2^ values, permutation test p-values (N=5000) and sample sizes for each network are provided in the inset table.

**Supplementary Fig. 5:**
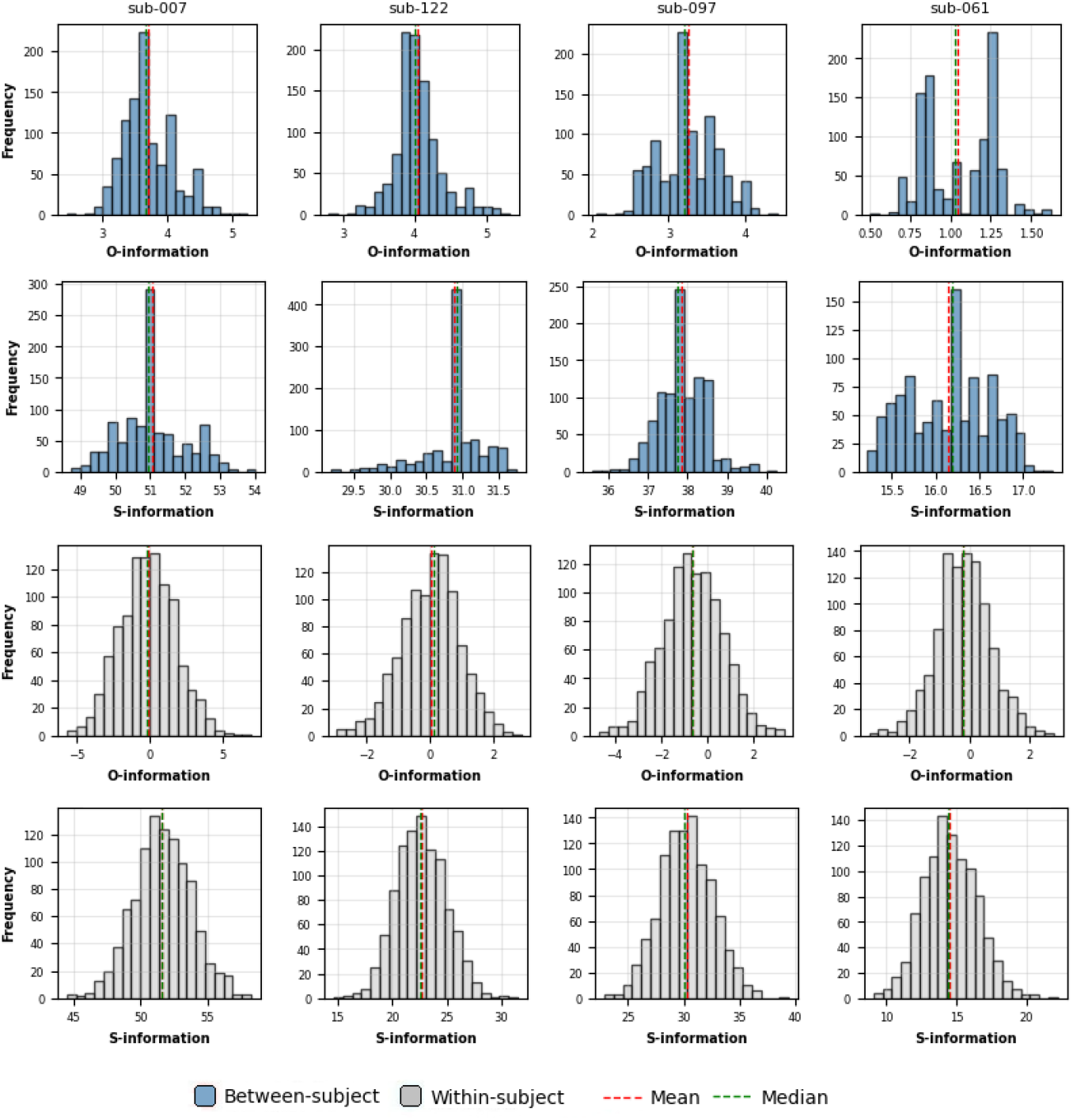
Z_b_-score and Z_w_-score reference distributions exhibit distinct visual differences. Histograms outlining the distribution of raw S- and O-information values for the EEN of a subset of patients with each column representing a single subject. Z_b_-score reference distributions are bootstrapped (1000 resamples with replacement) for visualisation. Null distributions appeared mostly Gaussian whereas distributions of matched healthy control networks appeared to potentially violate normality assumptions.

**Supplementary Fig. 6:**
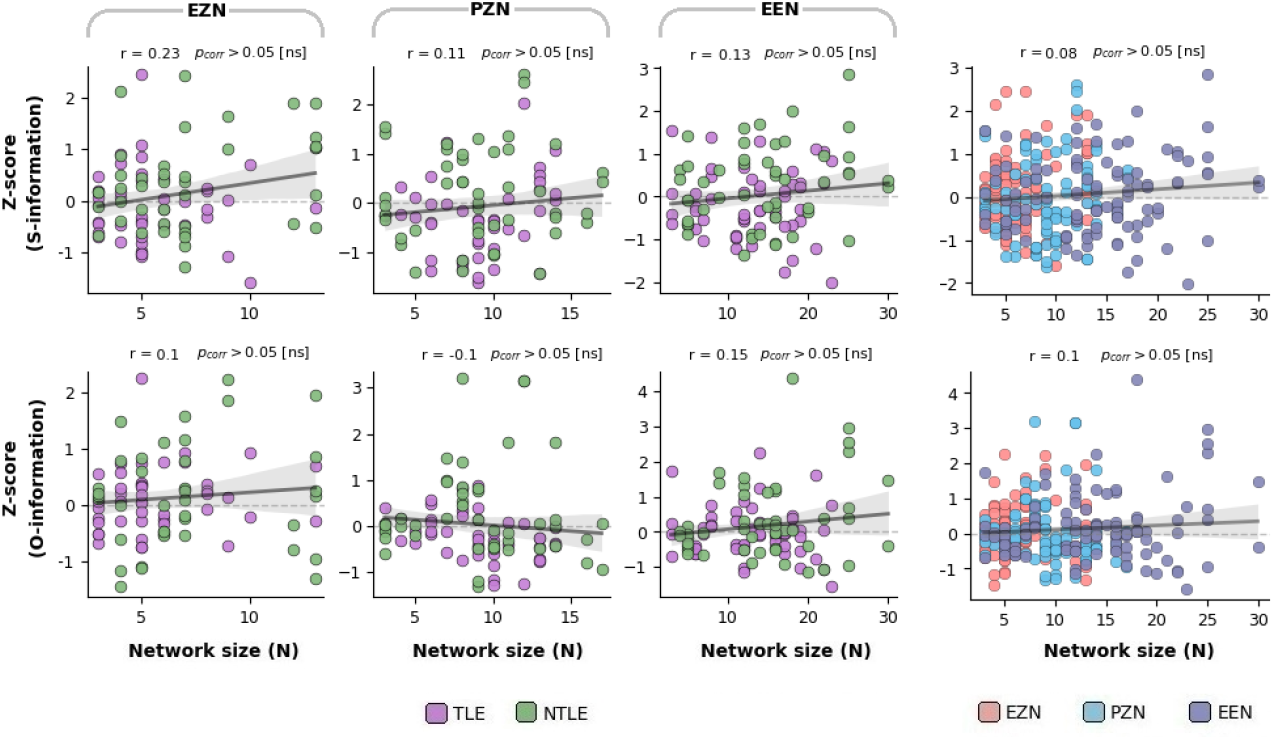
Z-scoring diminishes S- and O-information inflations with network size and supports comparison between heterogeneous patient groups. Scatter plots visualising the relationship between network size (number of nodes N) and both Z_b_- and Z_w_-scores for each network partition. Linear regression lines fitted across all subjects per network with 95% CI for regression computed via bootstrap (1000 iterations; grey shaded band). Grey dashed reference lines indicate deviations of zero.

## Notes

### Competing Interest Statement

The authors have declared no competing interest.

